# Establishment of Stable Immortalized Human Choroidal Melanocytes for Ocular Research

**DOI:** 10.64898/2026.02.16.706000

**Authors:** Aurélie Fuentes-Rodriguez, Andrew Mitchell, Vincent Gélinas, Kelly Coutant, Arnaud Droit, Solange Landreville

**Author notes:** **Corresponding Author:** Pr. Solange Landreville, Department of Ophthalmology and Otorhinolaryngology-Cervico-Facial Surgery, Hôpital du Saint-Sacrement, 1050 chemin Sainte-Foy, Room H2-02, Quebec City (QC), G1S 4L8, Canada.

## Abstract

**Purpose:** The short lifespan of primary normal choroidal melanocytes (NCMs) *in vitro* represents a major barrier to mechanistic, functional, and translational studies of choroid biology and uveal melanoma (UM). This study aimed to establish and characterize immortalized human NCM lines that retain melanocytic function, maintain a non-cancerous profile, and are amenable to gene editing.

**Methods:** NCMs from four donors were immortalized by lentiviral transduction of Cyclin-dependent kinase 4 (CDK4^R24C^), Cyclin D1, and human Telomerase reverse transcriptase (hTERT), establishing NCM-K4DT lines. Their morphology, melanocytic marker expression, proliferation and functional properties (melanin synthesis, tyrosinase activity) were evaluated. Genomic stability was assessed by targeted mutation profiling, karyotyping, and copy number variation analysis. The tumorigenicity was tested in immunodeficient mice. Plasmid-based CRISPR/Cas9 editing was performed to determine their suitability for gene editing.

**Results:** NCM-K4DT lines retained dendritic-shaped morphology, pigmentation, and expression of PMEL, TYRP1, Melan-A, and SOX10. Cells exhibited enhanced proliferative capacity with preserved cell cycle regulation. Melanin production and tyrosinase activity were comparable to primary NCMs. Genomic profiling confirmed the absence of UM-associated driver mutations and chromosomal abnormalities. *In vivo* growth assays demonstrated no tumorigenic potential. Notably, NCM-K4DT cells were efficiently edited by CRISPR/Cas9.

**Conclusions:** NCM-K4DT lines represent stable, non-cancerous, and genetically tractable models for studying choroidal melanocyte biology, modeling UM-associated mechanisms, and advancing therapeutic development in ocular research.

## Introduction

Normal choroidal melanocytes (NCMs) are the pigment-producing cells within the uveal tract (i.e., iris, ciliary body, choroid) and contribute among others to ocular light absorption, protection from photo-toxicity, and immunity.^1,2^ They are also implicated in ocular pathologies, including age-related macular degeneration, albinism, Waardenburg syndrome and uveal melanoma (UM), the most common primary intraocular malignancy in adults.^2–5^ Progress understanding NCM biology has been limited by the replicative senescence of primary cells cultured *in vitro* and the limited availability of donor tissues, which restrict the scale and reproducibility of mechanistic studies.^6^ Consequently, most studies rely on short-lived primary NCM cultures, non-choroidal melanocytes or established UM cell lines,^1,6–15^ each of which has important limitations for modeling early disease events and normal physiological function, and collectively hinder long-term genetic or pharmacological experiments.

Immortalization can overcome these constraints, but conventional approaches utilizing the SV40 large T antigen or human papilloma virus (HPV) E6/E7 oncoproteins disrupt tumor suppressor pathways (p53/pRb) and compromise genomic stability.^16,17^ We therefore adopted the K4DT immortalization method, which uses targeted genetic modifications (i.e., CD**<u>K4</u>**^R24C^, Cyclin **<u>D</u>**1, h**<u>T</u>**ERT) to modulate specific, downstream cell-cycle regulators and telomere maintenance, extending proliferative capacity while minimizing perturbations to tumor suppressor pathways and preserving genomic integrity.^18^

We first confirmed that the K4DT method successfully extends the proliferative capacity of NCMs, then demonstrated that these cells retain their melanocytic morphology and key functional characteristics. We further showed that NCM-K4DT lines maintain genomic stability, lack oncogenic driver mutations associated with UM, and do not exhibit tumorigenic behavior *in vitro* and *in vivo*. Finally, we have established NCM-K4DT as a versatile experimental platform for advanced genetic manipulation, including CRISPR-based gene editing, disease modeling and functional assay development.

## Materials and Methods

### Cell models

This research was approved by our Institutional Review Board (CHU de Québec-Université Laval Research Center; projects #2021-5273 and #2012-1483) and conducted in compliance with ARVO Best Practices for Using Human Eye Tissue in Research (Nov2021). Research-grade human eyeballs were provided by our local eye bank (Banque d’Yeux du Centre Universitaire d’Ophtalmologie, Quebec City, QC, Canada) and Héma-Québec (Quebec City, QC, Canada), with next-of-kin consent. Donor information is summarized in Table 1. Cells were isolated using a previously established method and cultured in a melanocyte growth medium, consisting in 1:1 mixture of DMEM (Corning) and Ham’s F12 medium (Wisent), supplemented with 10% FBS, 10 ng/mL cholera toxin (MilliporeSigma), 100 nM phorbol-12-myristate 13-acetate (MilliporeSigma), and penicillin/streptomycin (Corning).^6,7,10,15^ To select out only melanocytes from the pool of choroidal cells, 100 ug/mL geneticin G418 (Corning) was added to cultures from passages 0 to 5 only. For comparison and methodological validation, UM cell lines (Mel285, 92.1, Mel202, Mel270, Mel285, MP41, MP46, OMM2.3, UPMD1) and human embryonic kidney cells (HEK293T) were maintained under standard culture conditions.^19–25^ All cell types were grown at 37°C in a humidified atmosphere with 5% CO_2_ and were tested routinely for mycoplasma infection (MycoAlert^TM^ PLUS Detection Kit, Lonza).

**Table 1.** Information on NCM donors.

| Donor | Age /<br>sex | PPDI | Extraction<br>date (year) | Immortalization<br>passage | Cause of death |
| --- | --- | --- | --- | --- | --- |
| 1 | 51 / X | 1 | 2013 | P11 | Breast cancer |
| 2 | 66 / Y | 2 | 2021 | P7 | Lung cancer |
| 3 | 48 / Y | 4 | 2021 | P9 | Jaundice, weight loss |
| 4 | 48 / X | 3 | 2019 | P11 | Rectal cancer, liver metastases |

**PPDI**, postmortem delay between death and isolation; **P**, passage

### Plasmid construction

The pLEFS-CDK4-R24C-CyclinD1-Bleo lentiviral plasmid was generated using the pbabe-cyclinD1+CDK4R24C plasmid (a gift from Pr. Christopher Counter; Addgene plasmid #11129) and the pLEFS-AkaLuc-tagGFP2-BleoR plasmid (a gift from Dr. Darin Bloemberg, Novo Nordisk). The CDK4-R24C sequence was amplified by PCR, introducing a BamHI restriction site at the 5′ end and removing the stop codon while adding a NheI site at the 3′ end. The PCR product and the pLEFS-AkaLuc-tagGFP2-BleoR backbone was digested and ligated using the NEB Quick Ligation Kit (New England Biolabs), replacing the AkaLuc sequence with CDK4-R24C. Subsequently, tagGFP2 was replaced with Cyclin D1 by Gibson assembly using the Gibson Assembly Master Mix (New England Biolabs). To generate the third-generation pLV-hTERT-hygro plasmid, pLV-hTERT-IRES-hygro (a gift from Pr. Tobias Meyer; Addgene plasmid #85140) was digested with ClaI and XbaI and cloned into pLV-CMV-LoxP-DsRed-LoxP-eGFP (a gift from Pr. Jacco van Rheenen; Addgene plasmid #65726), which contains a chimeric 5′ LTR and 3′ SIN LTR. This produced an intermediate plasmid containing a puromycin resistance cassette (pLV-hTERT-puro, third-generation). To reintroduce the hygromycin resistance gene, the original second-generation pLV-hTERT-IRES-hygro plasmid was digested with KpnI, and the insert was ligated into pLV-hTERT-puro (third generation). Correct orientation of the KpnI insert was confirmed by Sanger sequencing. Plasmids were transformed into NEB Stable Competent E. coli (New England Biolabs, C3040H) and purified using the Qiagen EndoFree Plasmid Maxi Kit (Qiagen). All constructs were verified by Sanger sequencing (Sanger Sequencing Platform, CHU de QuébecUniversité Laval Research Center). A list of Gibson assembly primers, sequencing primers, and graphical cloning maps is provided in the Supplementary Data. For initial proof-of-concept experiments, pHAGE-CDK4-R24L (a gift from Gordon Mills & Kenneth Scott; Addgene plasmid #116217) was used as a readily available lentiviral vector encoding a p16^INK4A^-resistant CDK4 allele. Following confirmation of functional efficacy in pilot experiments (Supplementary Fig. S1), a custom pLEFS-CDK4-R24C-CyclinD1-Bleo construct was generated incorporating the R24C mutation, consistent with previously published K4DT immortalization strategies.

### Lentiviral production

Lentiviral particles were produced using a third-generation packaging system in HEK293T cells (ATCC, #CRL-3216). Cells were seeded in 10-cm dishes at 70–80% confluency 24 hours prior to transfection. Transfer plasmids encoding CDK4–CyclinD1 or hTERT were co-transfected with the ViraPower Lentiviral Expression System (Thermo Fisher Scientific) at a molar ratio of 2:1:1:1 (transfer:P1:P2:VSVG) using PEI MAX 40K (Polysciences), following the manufacturer’s instructions. The transfection mixture was prepared in Opti-MEM (Gibco), incubated for 15 minutes, and then added to the cells. After 6-8 hours, the medium was replaced with DMEM supplemented with 10% FBS, 1% non-essential amino acids, 10 mM HEPES, and 1 mM sodium butyrate. At 72 hours post-transfection, supernatants containing lentiviral particles were collected, centrifuged at 3,000 rpm to remove debris, filtered through a 0.45-µm membrane, and concentrated by ultracentrifugation using an SW 32 Ti rotor at 14,000 rpm for 4 hours at 4°C. Viral pellets were resuspended in PBS overnight at 4°C, centrifuged at 7,000 × *g* for 5 minutes to remove aggregates, and stored at -80°C. Viral titers were estimated using HEK293T cells.

### Generation of CDK4/CyclinD1 and hTERT (K4DT)-expressing NCMs

Initial pilot experiments demonstrated that transgenic expression of either CDK4-Mut, Cyclin D1 or hTERT alone was insufficient for immortalization (Supplementary Fig. S1). This is consistent with a previous study in cutaneous melanocytes.^26^ In contrast, incorporation of all K4DT transgenes into a single tri-cistronic lentiviral vector was impractical due to the resulting large vector size and the inherent challenges associated with hTERT expression (long coding sequence, GC-rich content, and propensity for recombination), which would be expected to substantially reduce viral titre. Therefore, a two-step sequential transduction strategy was used. First, primary NCMs (80,000 cells/well) were seeded in 12-well plates. After 72 hours, the medium was replaced with NCM medium containing 5 µg/mL polybrene (Millipore-Sigma) and CDK4/CyclinD1 lentiviral particles at an MOI of 5. After 8 hours, the virus-containing medium was replaced with fresh NCM medium. Three days later, cells were split into medium containing 500 µg/mL Zeocin (InvivoGen) and selected for 12 days. Next, zeocin-resistant CDK4/CyclinD1-expressing NCMs were seeded in 6-well plates (80,000 cells/well) for 72 hours and then transduced with hTERT lentivirus in NCM medium containing 5 µg/mL polybrene at an MOI of 5. After 8 hours, the medium was replaced. Three days later, cells were transferred into medium containing 150 µg/mL hygromycin (Wisent) and selected for 10 days. Passage numbers for the resulting K4DT cell line reflect the total passages since initial isolation from the donor.

### Generation of Luc2/tdTomato-expressing cell lines

Luciferase/fluorescent reporter lentivirus was produced as described above using the pCDH-EF1-Luc2-P2A-tdTomato transfer plasmid (a gift from Pr. Kazuhiro Oka; Addgene plasmid #72486). NCM-K4DT cells or 92.1 UM cells were infected in the presence of 5 µg/mL polybrene at an MOI of 3. Two rounds of fluorescence-activated cell sorting (BD FACSMelody™, BD Biosciences) were performed to obtain pure, homogeneous fluorescent populations. Although we were initially successful in transducing primary NCMs, we were unable to obtain a stable, fully pure population of Luc2/tdTomato-expressing cells before they became senescent.

Luciferase expression was confirmed using a plate-based bioluminescent assay. Briefly, cells were lysed in 1X radioimmunoprecipitation assay (RIPA) buffer (50 mM Tris-HCl, pH 7.4, 150 mM NaCl, 0.6% NP-40; 5×10^6^ cells per mL) on ice for 30 minutes, before being centrifuged for 20 minutes at 16,000 x *g* to remove debris. The supernatant was removed and diluted 5X in distilled H_2_O and then subsequently serially diluted. Near 98 µL of cell supernatant were added to each well of a white 96-well plate (Sarstedt), followed by 100 µL of TMCA buffer (5 mM MgCl_2_ (Millipore Sigma), 0.25 mM CoA (GoldBio), 0.15 mM ATP (BioBasic), 100 mM Tris-HCl (Millipore Sigma) pH 7.8). Finally, 2 µL of luciferin stock solution (15 mg/mL of D-luciferin potassium salt (GoldBio) in biology grade H_2_O) was quickly added to each well and the bioluminescence was immediately detected on an iD5 SpectraMax microplate reader (Molecular Devices).

### Polymerase chain reaction (PCR)

To confirm the insertion of the lentiviral constructs into the host genome, PCR was performed to specifically detect the plasmid DNA. Primers were designed to overlap the promoter and coding sequences to ensure the transgene and not the native genes were being detected. Since CDK4^R24C^ and Cyclin D1 were in the same plasmid, only one set of primers was needed. Genomic DNA was extracted from cells using the DNeasy Blood and Tissue Kit (Qiagen), and 500 ng of DNA was used as template for PCR amplification using the Q5 High-Fidelity 2X Master Mix (New England Biolabs) with gene-specific primers (Table 2). Amplification was performed with an initial denaturation step at 98°C for 60 seconds, followed by 30 cycles: denaturation at 98°C for 10 seconds, annealing at 68°C for 20 seconds, and extension at 72°C for 20 seconds. Once the cycles were completed, a final extension at 72°C for 2 minutes was done. PCR products were resolved on a 3.5% agarose gel and electrophoresed for 60 minutes at 100 V. Bands were visualized under UV illumination (Azure Biosystems c150) to confirm the presence of the expected amplicons corresponding to the inserted genes.

**Table 2.**
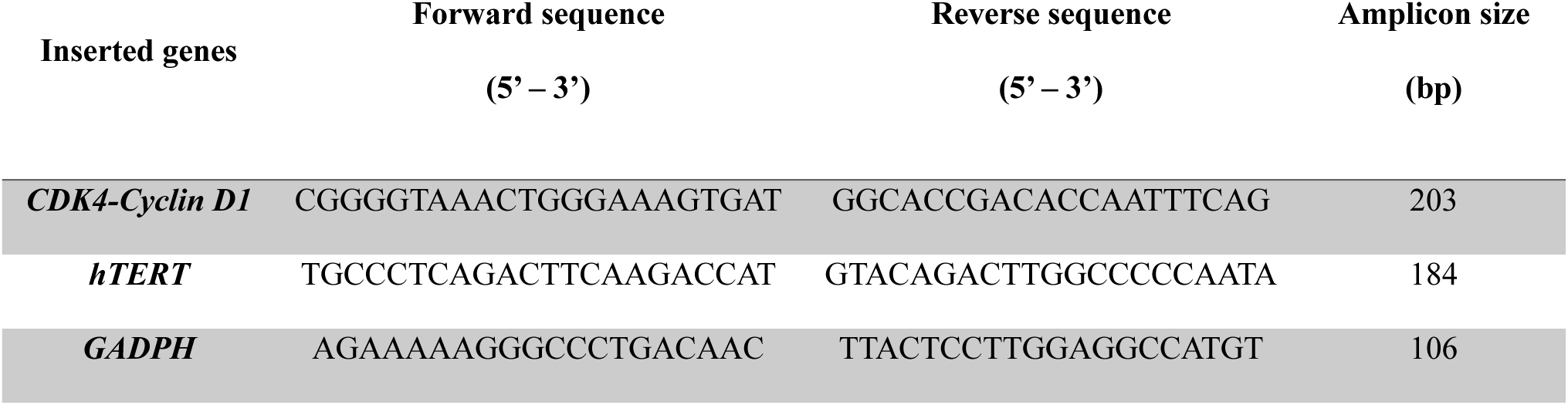
PCR primer sequences.

### Western blotting

Cell pellets from NCM-K4DT lines and primary culture controls were lysed in RIPA buffer, and the total protein concentration was determined with the Pierce^TM^ BCA Protein Assay Kit (Thermo Fisher Scientific).^13^ Twenty µg of protein per sample were loaded onto a 4-15% Mini-PROTEAN TGX Precast Protein Gels (Bio-Rad). Proteins were separated by electrophoresis and transferred onto nitrocellulose membranes. Following the transfer, membranes were blocked in TBS-T (TBS + 0.1% Tween-20) containing 5% non-fat dry milk for 1 hour at room temperature (RT) and subsequently incubated overnight at 4°C with primary antibodies CDK4 (1:1,000; Cell Signaling, #12790), Cyclin D1 (1:1,000; Cell Signaling, #2978), hTERT (1:1,000; Abcam, #32020) or p21 (1:500; Santa Cruz Biotechnology, #6246). The next day, membranes were incubated for 1 hour at RT with a pan-Actin antibody (1:10,000; Cedarlane, #CLT9001), then washed in TBS-T and incubated with the appropriate HRP-conjugated secondary antibodies: anti-mouse IgG or anti-rabbit IgG (1:2,500; Jackson Immuno Labs, #115-036-003 and #313-036-003) at RT for 1 hour. Chemiluminescence was visualized with the WesternSure^TM^ PREMIUM Substrate and the C-DiGit Blot Scanner (LI-COR).

### Immunofluorescence

Cells (50,000 cells) were seeded on coverslips and incubated at 37°C and 5% CO_2_ for 48 hours, then fixed with 4% paraformaldehyde, permeabilized with 0.1% Triton X-100 and blocked with 4% bovine serum albumin (BSA) at RT. Cells were incubated with primary antibodies against Ki67 (1:100; BD Biosciences, #556003; proliferation marker), TYRP1 (1:1,000; Santa Cruz Biotechnologies, #25543; melanocyte marker), PMEL (1:50; Abcam, #137078; melanocyte marker) or SOX10 (1:100; Abcam, #218522; melanocyte marker), then washed and incubated 1 hour at RT with Alexa Fluor-594 conjugated secondary antibodies (1:250; Cell Signaling Technology, #A21442 and #A21203) along with Phalloidin 488 (1:250; Cayman Chemical, #20549; F-actin marker) and DAPI (1:2,000; Thermo Fischer Scientific, #D3571; nuclear marker). Coverslips were washed, mounted with Fluoromount-G (Southern Bio), and stored at 4°C overnight. Images were acquired using a Zeiss LSM800 confocal microscope with identical settings across conditions. For quantification, images were processed in ImageJ (NIH, https://imagej.net/ij/). Nuclei were segmented using the DAPI channel as a reference, and regions of interest (ROIs) were defined for each nucleus. The proportion of Ki67-positive nuclei was calculated by measuring the overlap between the nuclear ROIs and Ki67 fluorescence signal, using a consistent intensity threshold across samples. At least five random fields per condition were analyzed, and data were expressed as the percentage of Ki67-positive cells relative to the total number of nuclei.

### Population doubling and cell cycle analysis

Population doubling (PD) was calculated by monitoring the growth of NCM-K4DT low- (LowP; passages 16-20), high-passage (HighP; passage 25), and control cells (passages 9-13) over 14 consecutive days, with counts taken at 72- to 96-hour intervals. PD was calculated using the formula: PD = x + 3.322*(log_10_*N_f_* – log_10_*N_i_*) where x is the cumulative PD from the previous passage, *N_f_* is the final cell number after trypsinization, and *N_i_* is the initial number of cells plated. To further evaluate the cell cycle progression, NCM-K4DT lines (LowP and HighP) and controls were fixed in 70% ethanol, treated with 100 µg/mL RNase A, stained with 50 µg/mL propidium iodide (PI), and analyzed by flow cytometry (BD Accuri C6 Plus). Gating strategies were applied to exclude doublets and cell debris, ensuring accurate quantification of G0/G1, S, and G2/M phases.

### Melanin quantification

Cells were collected (2×10^5^), washed twice with PBS and centrifugated at 12,000 × *g* for 10 min. Pellets were solubilized in 100 μL οf 1N NaOH-10% DMSO at 80°C for 1 hour, and the melanin content was determined by measuring the absorbance at 405 nm with a normalization performed with a BCA quantification post-measurement.^27^ A standard series of known BCA concentrations was prepared in separate wells, and 100 µL of prepared BCA solution were added to each well. Plates were incubated at 37°C for 30 minutes, and the absorbance was recorded at 562 nm for protein normalization.

### L-DOPA tyrosinase activity assay

To assess L-DOPA tyrosinase activity, NCM-K4DT lines and primary culture controls were seeded at a density of 2.5×10^4^ cells per well in a 96-well clear-bottom microplate, in triplicates. Controls included a blank (no cells) and a negative control (choroidal fibroblasts). Cells were incubated for 24 hours at 37°C with 5% CO_2_ before measuring the DOPA reaction. On the day of the assay, the medium was removed, and cells were gently washed with PBS. Freshly prepared 1 mM L-DOPA (Thermo Fisher Scientific) was then added (100 µL per well), and absorbance at 475 nm was monitored at multiple time points (T0 and every 15 minutes) to track dopachrome formation.^28^ A time-zero measurement was performed to establish a baseline and determine the rate of dopachrome formation, with normalization on total protein. The protein extraction was performed by removing the L-DOPA solution, washing the wells with PBS, and lysing the cells with 100 µL of RIPA buffer for 10 minutes at RT on an orbital shaker. Total proteins were quantified with a BCA assay as described above.

### Genotyping

To assess the mutational status of key genes associated to UM, specific hotspot mutations in *GNAQ* (Q209), *GNA11* (Q209), *CYSLTR2* (L129), *PLCB4* (D630), *SF3B1* (R625), and *EIF1AX* (G5/6/9) genes were screened using specific primers (Table 3). Genomic DNA was extracted from NCM-K4DT, primary culture controls and eight UM cell lines using a standard QuickExtract (Lucigen), PCR amplification was performed as previously described. Final PCR products were sent to the Sanger Sequencing Platform (CHU de Québec-Université Laval Research Center) for purification and sequencing. Results were compared to known reference sequences on Benchling (https://benchling.com).

**Table 3.**
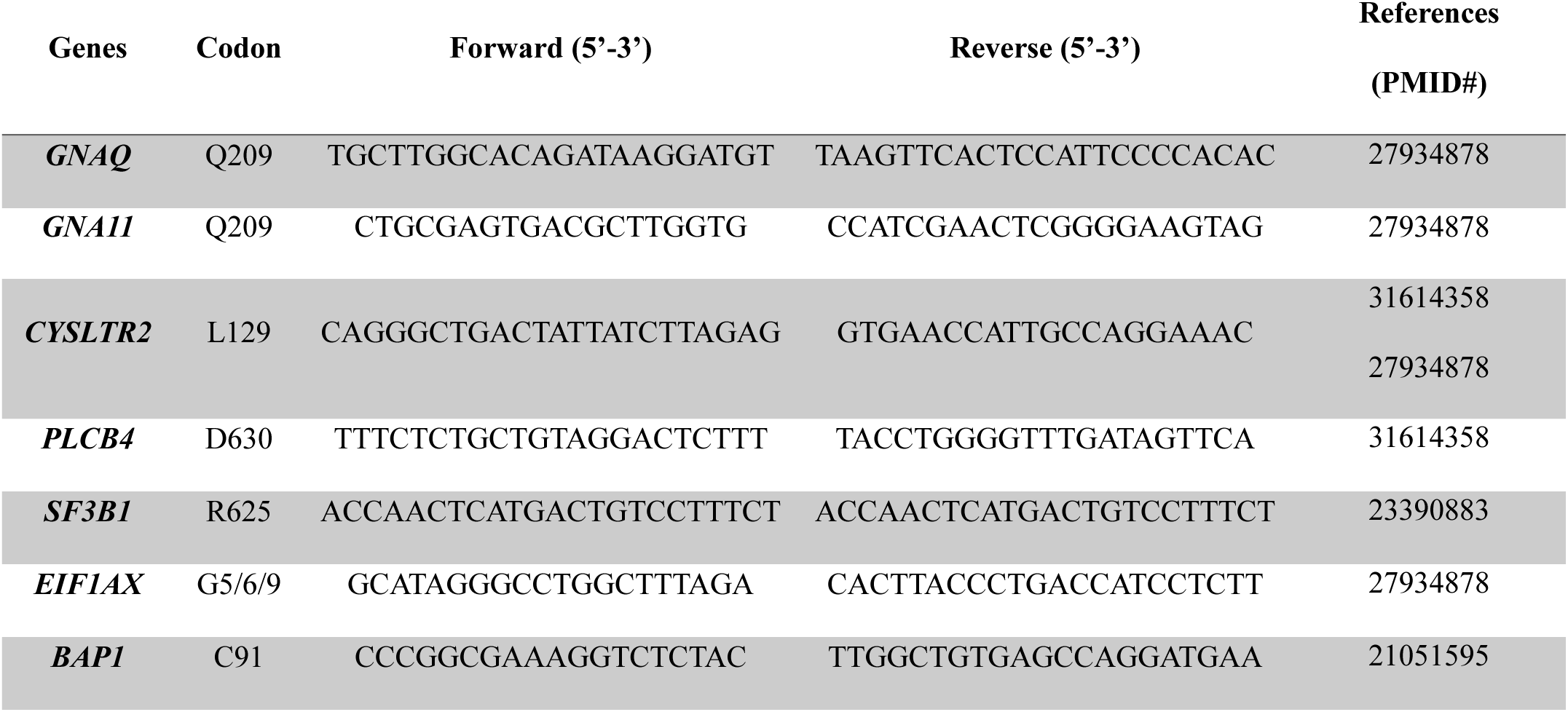
Genotyping primer sequences.

### Karyotyping and copy number variation (CNV) analysis

Frozen cell pellets were submitted to Genome Québec (Montreal, Canada) for whole-genome sequencing–based karyotyping using the Illumina Infinium Global Diversity Array (GDA-8 v1.0). Genotype intensity data were exported from GenomeStudio containing per-marker Log R Ratio (LRR) and B Allele Frequency (BAF). Processing was performed in R (packages: data.table, dplyr, ggplot2, DNAcopy). For each sample, 0.5–99.5% tails were winsorized; when enabled, per-bin GC% (GRCh37/hg19) was used for LOESS GC correction after clamping bin ends to chromosome boundaries, following established SNP-array wave/GC adjustments.^29^ For binning and segmentation, probes were aggregated to 200 kb bins (median per bin; bins with <3 probes dropped) and the denoised bin-level LRR were segmented with circular binary segmentation (DNAcopy; α=0.01; SD=1.0, min.width=5), a standard approach for array CGH/SNP data.^30^ Sex was inferred from chrX heterozygosity (BAF) and chrY LRR; PAR1/2 were treated as autosomal, and bin LRR were converted to copy number as CN_estimate_=CN_expected_ x 2^LRR^ with CN_expected_=2 for autosomes/PARs; for XX: *X*_non-PAR_=2, *Y*_non-PAR_=0; for XY: *X*_non-PAR_=1, *Y*_non-PAR_=1.^31^ Genome-wide RD[CN] plots were generated with segment overlays and exported per-bin and per-segment tables; QC included autosomal LRR SD after cleaning, retained-bin counts, and concordance of the sex call with chrX/chrY signals.

### In vivo tumorigenicity assessment

All animal experiments were conducted in voluntary compliance with the ARVO Statement for the Use of Animals in Ophthalmic and Vision Research and were approved by our institutional animal experimentation committee (Université Laval; protocol #2023-1356). One million cells of 92.1-Luc2-tdTomato or NCM-K4DT-Luc2-tdTomato suspended in a 1:1 mixture of PBS and Matrigel^®^ (Corning) were injected subcutaneously into the left and right flanks of immunodeficient mice (NOD-Prkdc^em26Cd52^Il2rg^em26Cd22^/NjuCrl; Charles River Laboratories, #572). Animals were monitored weekly for tumor formation, overall health, and signs of distress in accordance with institutional animal care guidelines. Tumor growth was measured with calipers, and tumor volume was calculated as ½×length×width^2^. In addition, longitudinal tumor progression was monitored at 2-, 4-, 8- and 12-weeks post-injection using bioluminescence with the IVIS Lumina II System (PerkinElmer). For imaging, mice were anesthetized with isoflurane and received an intraperitoneal injection of D-luciferin potassium salt (GoldBio; 15 mg/mL in D-PBS (Wisent)) at a dose of 100 µL per 10 g of body weight. Animals were imaged 11 minutes following injection, corresponding to the empirically predetermined peak luminescent signal. The bioluminescent signal was quantified as total photon flux (photons/s) using the Living Image software (PerkinElmer). Animals were euthanized after 12 weeks, or earlier if tumors exceeded the maximum allowable volume (2 cm^3^). Following euthanasia, tumors or Matrigel^®^ plugs were excised, imaged for fluorescence or luminescent signal and fixed in 4% paraformaldehyde solution in PBS.

### SCIP-based CRISPR editing

To generate targeted mutations in NCM-K4DT lines, we employed a SCIP system (self-cleaving integrating plasmid) for CRISPR-Cas9-mediated gene editing using the pQCi plasmid (a gift from Dr. Darin Bloemberg; Addgene #154090).^32^ CRISPR single-guide RNAs (sgRNAs; Integrated DNA Technologies) and homology arms (HAs; Twist Bioscience) harboring UM mutations *GNAQ^Q^*^*209*^*^L^*, *GNA11^Q^*^*209*^*^L^*and *BAP^C91G^* were designed using Benchling and cloned following the pQCi backbone protocol and SCIP technique.^32^ A silent mutation was introduced within the PAM sequence in the HAs to prevent re-cutting of the edited locus by Cas9 following HDR. This synonymous change preserves the protein sequence while protecting the correctly edited allele from repeated cleavage and unintended indel formation. Plasmid DNA was amplified in DH5α E. coli and purified using NucleoSpin® Plasmid Transfection-grade kit (Machery-Nagel). NCM-K4DT cells were seeded at 1.7×10^4^ cells per cm^2^ in 100 mm dish two days prior. Transfections were performed using a Neon Invitrogen electroporator as previously described,^6^ with 500 ng of SCIP per condition in triplicates. Twenty-four hours post-transfection, cells were stained with the Ghost Dye Red 780 Fixable Viability Dye (Tonbo Biosciences) and sorted using a FACSMelody^TM^Cell Sorter (BD Biosciences) to enrich for viable, EGFP-expressing populations. The sorted cell pools were expanded for two passages and subsequently analyzed. Editing efficiency and indel spectra were quantified from Sanger chromatograms using Inference of CRISPR Edits (ICE; Synthego) with default parameters, by comparing each edited sample to its corresponding wild-type control trace.^33^ In parallel, Mel285 and HEK293T cells were processed under identical conditions to validate the transfection and sorting flow.

### Statistical analyses

All statistical analyses were performed using GraphPad Prism (version 10.6.1, GraphPad Software). Data are presented as mean ± standard error of the mean (SEM). Comparisons between two groups were performed using unpaired two-tailed Student’s *t*-tests. For multiple group comparisons, one-way or two-way ANOVA followed by appropriate post hoc tests was used. A *p*-value < 0.05 was considered statistically significant.

## Results

### Generation of NCM-K4DT lines

To establish stable human immortalized choroidal melanocyte cell lines, we utilized a lentiviral transduction approach incorporating hTERT, CDK4^R24C^ and, Cyclin D1 (Fig. 1A). These constructs were designed to enhance cell cycle progression and extend replicative potential while preserving genomic integrity.^18^ The successful gene integration was confirmed by PCR (Fig. 1B) and Western blot analysis (Fig. 1C), demonstrating a statistically significant increase in hTERT (*p* < 0.001), CDK4 (*p* < 0.01), and Cyclin D1 (*p* < 0.05) levels in NCM-K4DT lines compared to primary culture controls (Fig. 1D).

**Figure 1.**
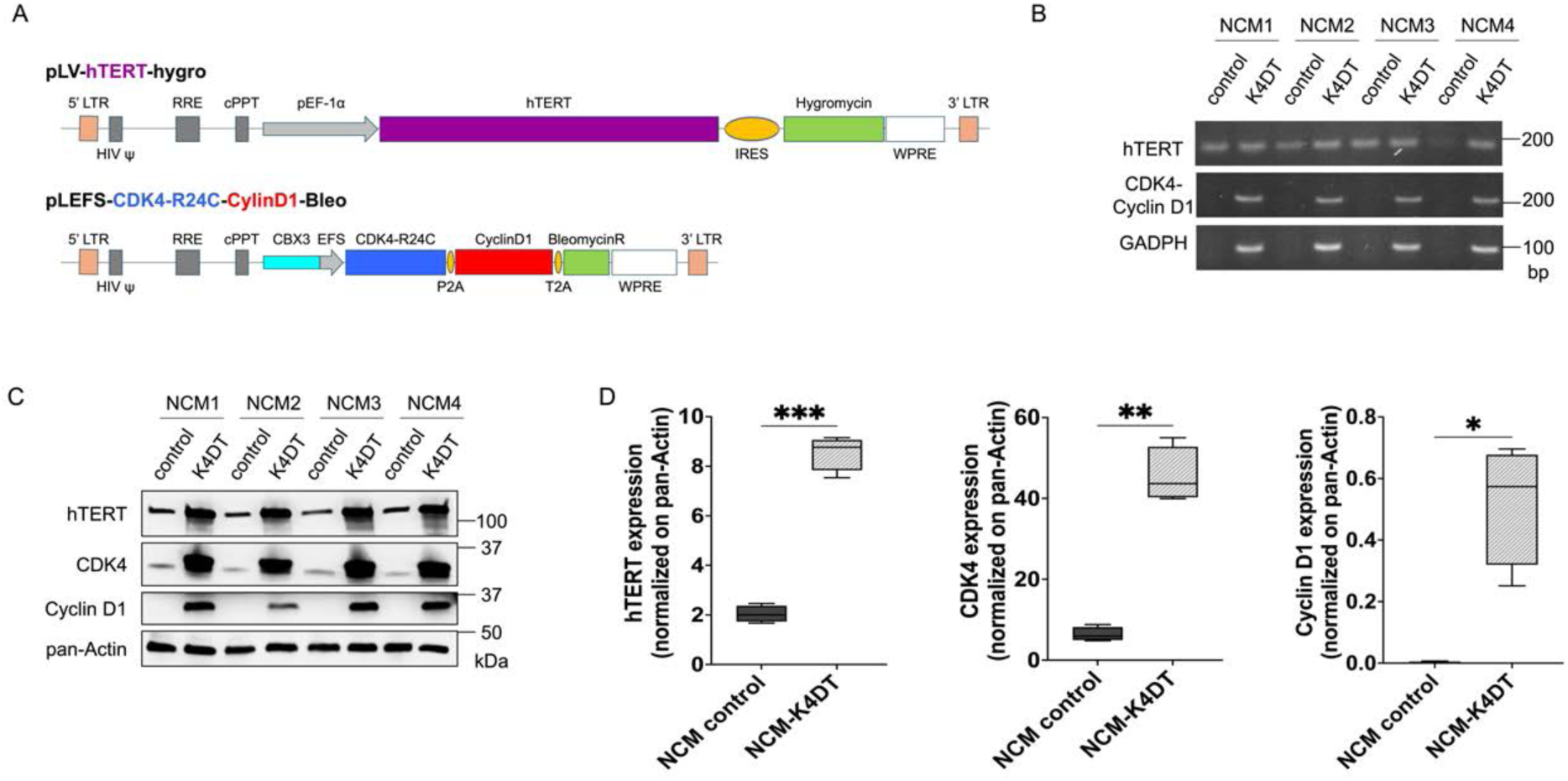
Generation and validation of NCM-K4DT lines expressing hTERT, CDK4^R24C^, and Cyclin D1 transgenes. (A) Schematic representation of the lentiviral vectors used for the expression of hTERT, CDK4^R24C^, and Cyclin D1 in primary NCMs. (B) PCR analyses confirming the insertion of hTERT, CDK4^R24C^, and Cyclin D1 transgenes in NCM-K4DT lines compared to controls. GAPDH served as loading control. (C) Western blot validation of transgene expression in immortalized NCM-K4DT cells, showing a robust expression of hTERT, CDK4, and Cyclin D1 compared to controls; pan-Actin served as loading control. (D) Densitometry analysis of protein expression levels demonstrating a significant upregulation of hTERT (\*\*\**p* < 0.001), CDK4 (\*\**p* < 0.01), and Cyclin D1 (\**p* < 0.05) in NCM-K4DT lines relative to primary cells.

### Preservation of melanocytic identity in NCM-K4DT cells

To confirm that NCM-K4DT lines retained key melanocytic properties, we evaluated the expression of melanocyte-specific markers (i.e., PMEL, TYRP1, Melan-A, SOX10). The Western blot analysis detected robust expression of PMEL, TYRP1 and Melan-A (Fig. 2A), and no significant difference compared to NCM controls (Fig. 2B). Immunofluorescence analyses further confirmed the expression of melanocytic markers PMEL, TYRP1, and SOX10 (Fig. 2C), indicating that their immortalization did not significantly alter the melanocytic phenotype.

**Figure 2.**
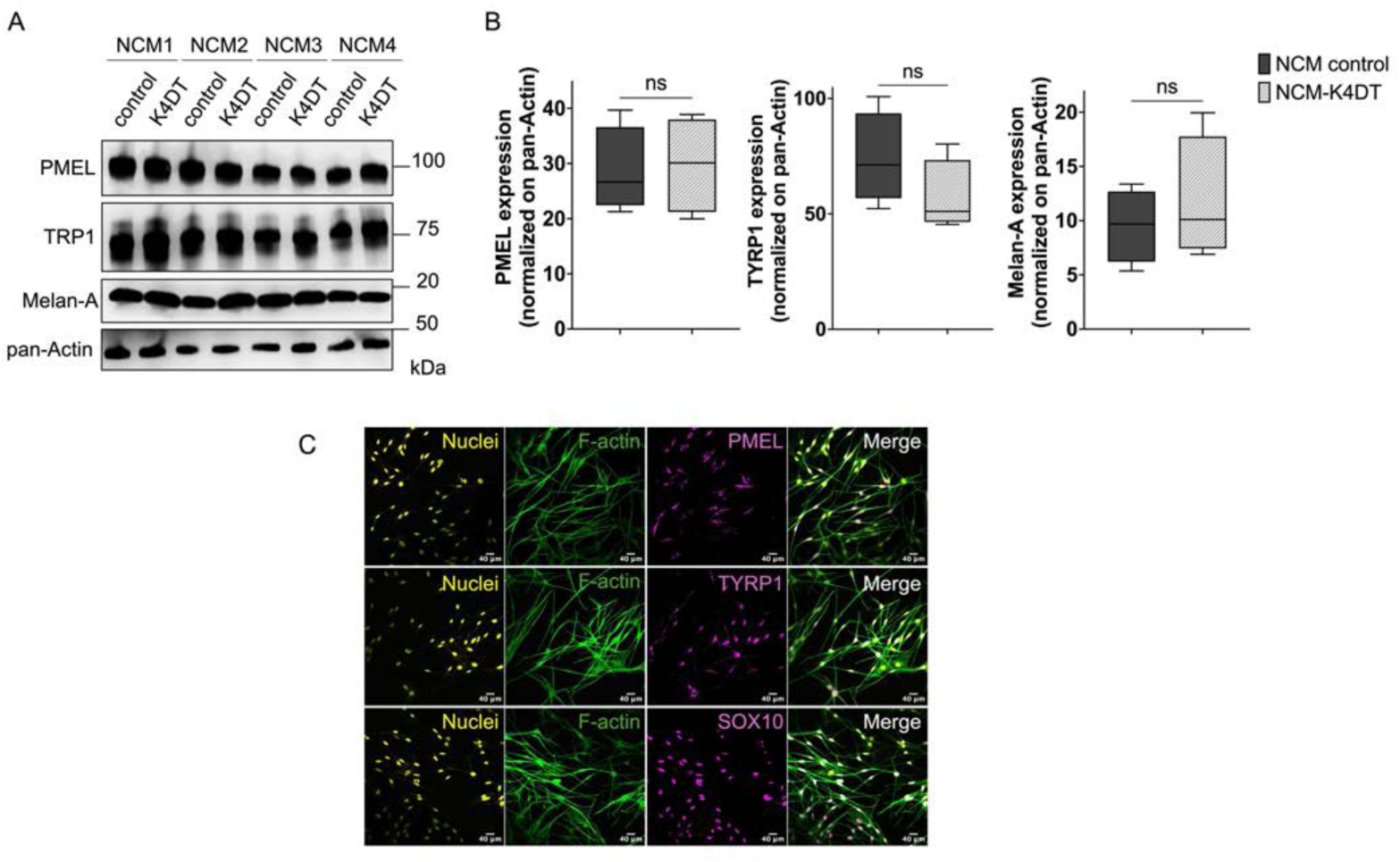
Preservation of melanocytic marker expression in NCM-K4DT lines. (A) Western blot analysis showing expression of melanocytic markers PMEL, TYRP1, and Melan-A in primary NCMs and NCM-K4DT lines, with pan-Actin as loading control. (B) Densitometry analysis of PMEL, TYRP1, and Melan-A protein expression levels, showing no significant differences between NCM controls and K4DT lines. (C) Immunofluorescence staining confirming the expression of PMEL, TYRP1, and SOX10 (purple) in NCM-K4DT lines; nuclei are stained with DAPI (yellow) and F-actin with phalloidin (green).

### Enhanced proliferation and preserved cell cycle regulation in NCM-K4DT lines

Initial pilot experiments were performed to determine whether expression of individual immortalization components was sufficient to sustain long-term proliferation of primary NCMs. Cells transduced with CDK4, CyclinD1, or hTERT alone showed reduced proliferative capacity compared to cells expressing all three factors (Supplementary Fig. S1A-B). In extended culture experiments spanning over 70 days, non-transduced NCMs exhibited a clear plateau in cumulative population doublings, consistent with replicative senescence, whereas cells expressing CDK4, CyclinD1, and hTERT continued to proliferate without an apparent decline in division rate (Supplementary Fig. S1C). These findings informed the final experimental design. Based on these pilot studies, the CDK4 and CyclinD1 transgenes were subsequently combined into a single lentiviral construct, while the hTERT transgene was delivered separately.

Using this optimized system, we assessed the proliferative advantage conferred by immortalization over 13 days (Fig. 3A). NCM-K4DT lines displayed a significantly faster proliferation rate compared to the primary NCMs (*p* < 0.0001), with notably shorter population doublings. To assess whether this proliferative capacity was maintained over extended culture, we compered the growth rate of early-passage (LowP) and late-passage (HighP) NCM-K4DT cells. Population doublings remained consistent between LowP and HighP populations, demonstrating that NCM-K4DT lines sustain stable, long-term proliferative capacity without evidence of senescence or growth arrest. Next, we performed cell cycle analysis to determine the distribution of immortalized melanocytes across cell cycle phases (Fig. 3B-C). Early- and late-passage NCM-K4DT cells exhibited a comparable distribution of cells in G1, S, and G2 phases compared to primary NCMs, suggesting that the immortalization process did not disrupt normal cell cycle regulation (Fig. 3B). However, compared to primary NCMs, NCM-K4DT lines showed a decreased percentage of cells in G0/G1 (*p* < 0.05) and an increased proportion in S and G2/M phases (*p* < 0.05 both phases for LowP; only S phase for HighP) (Fig. 3C). This shift toward active cell cycle phases is consistent with the shorter population doubling time observed in K4DT-immortalized cells

**Figure 3.**
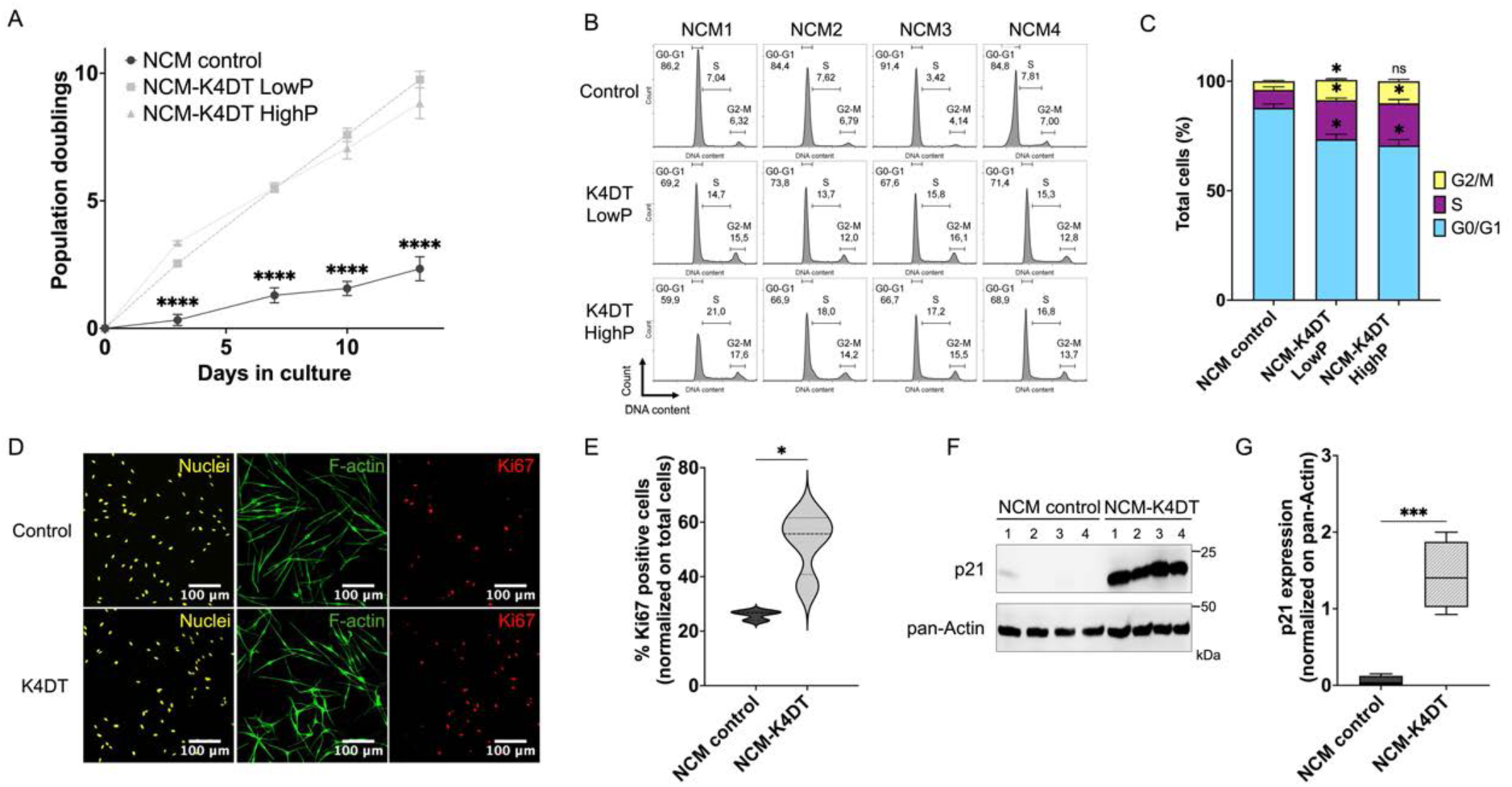
Enhanced proliferative capacity and cell cycle progression in NCM-K4DT lines. (A) Growth curve analysis showing significantly higher proliferation of NCM-K4DT lines compared to primary NCMs over 13 days (\*\*\*\**p* < 0.0001). (B) Plots illustrating cell cycle profiles in NCM controls and K4DT LowP/HighP lines and (C) cell cycle distribution determined by flow cytometry (\**p* < 0.05; ns: non-significant). (D) Representative immunofluorescence staining for Ki67 (red) in NCM2 cells; nuclei are counterstained with DAPI (yellow) and F-actin with phalloidin (green). (E) Quantification of Ki67-positive cells (\**p* < 0.05). (F) Representative Western blot showing p21 expression in NCM controls and K4DT cells. (G) Densitometry analysis of p21 expression levels normalized to pan-Actin (\*\*\**p* < 0.001).

Immunostaining for the proliferating marker Ki67 further validated the enhanced proliferative capacity of NCM-K4DT cells (Fig. 3D). The significantly higher proportion of Ki67-positive cells (*p* < 0.05) in immortalized melanocytes supports their sustained proliferative activity (Fig. 3E). Given that enforced expression of Cyclin D1 and CDK4 directly promotes cell cycle progression and enhanced proliferation, we next examined whether activation of endogenous cell-cycle inhibitory mechanisms accompanied the enhanced proliferative state of NCM-K4DT cells (Fig. 3F-G). Western blot analysis revealed a marked upregulation of the CDK inhibitor p21^34^ in NCM-K4DT lines relative to primary NCMs (*p* < 0.001), consistent with engagement of a compensatory checkpoint response that preserves downstream cell cycle control and restrains unchecked proliferation despite sustained mitogenic signaling.

Collectively, these findings confirm that NCM-K4DT lines maintain a robust yet controlled proliferative capacity, supporting their utility as a renewable *in vitro* model for choroidal melanocyte research.

### Preserved melanin synthesis in NCM-K4DT lines

The melanin content was qualitatively analyzed by visual comparison of cell pellets, where NCM-K4DT cells exhibited pigmentation in comparison to the unpigmented UM cell line Mel285 (Fig. 4A). Quantification of melanin levels revealed no significant difference between NCM-K4DT LowP or HighP and controls, suggesting that their immortalization did not impair melanin synthesis, even though K4DT cells showed a trend toward lower melanin levels (Fig. 4B). Given that tyrosinase activity assessed by the DOPA reaction (Fig. 4C), and melanosome marker expression (Fig. 2) were comparable between cell lines, we hypothesized that the increased proliferation rate of K4DT cells, rather than any defect in melanin biosynthesis, reduced melanin accumulation. To test this hypothesis, we reduced the concentration of PMA and cholera toxin by half in the culture medium to slow cell division and indeed observed an increase in melanin production (Supplementary Fig. S2C).

**Figure 4.**
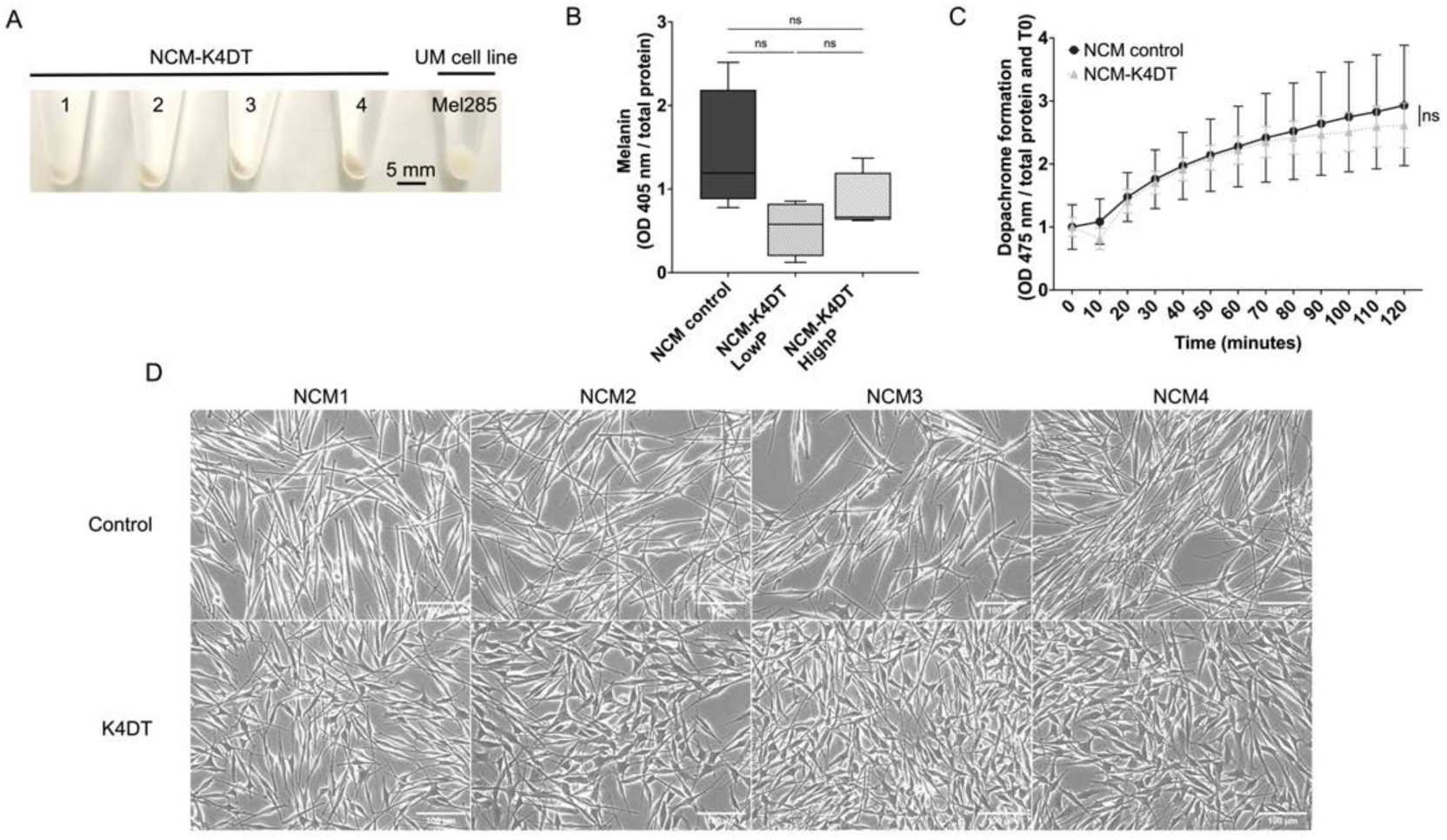
Melanocytic function assessment of NCM-K4DT lines. (A) Pigmentation in cell pellets obtained from NCM-K4DT lines and UM cells (Mel285). (B) Quantification of melanin content in NCM-K4DT lines compared to primary NCMs (ns: non-significant). (C) The dopachrome formation assay measuring the tyrosinase activity revealed comparable enzymatic function between NCM-K4DT lines and controls (ns: non-significant). (D) Phase-contrast microscopy images of NCM-K4DT cells showing a dendritic shape typical for melanocytes.

Additionally, the NCM-K4DT lines (Fig. 4D, lower panel) maintained a dendritic-shaped morphology characteristic of primary cultures of melanocytes (Fig. 4D, upper panel), further supporting their melanocytic identity.

### Genetic and chromosomal stability of NCM-K4DT lines

To ensure that NCM-K4DT lines retained a non-cancerous genomic profile, we performed mutation screening, chromosomal stability analysis, and *in vivo* tumorigenic assays. Recurrent mutations in UM include *GNAQ*, *GNA11*, *CYSLTR2*, *PLCB4*, *SF3B1*, and *EIF1AX* genes.^35^ To ensure that the rapid dividing nature of the K4DT cells did not increase their susceptibility to gain these mutations, we tested and compared the K4DT cells to UM cell lines with known mutations (Fig. 5A). These mutations were not present in the original donors and were not introduced after K4DT immortalization (Fig. 5A), in contrast to UM cell lines in which we were able to detect these common mutations as expected.

**Figure 5.**
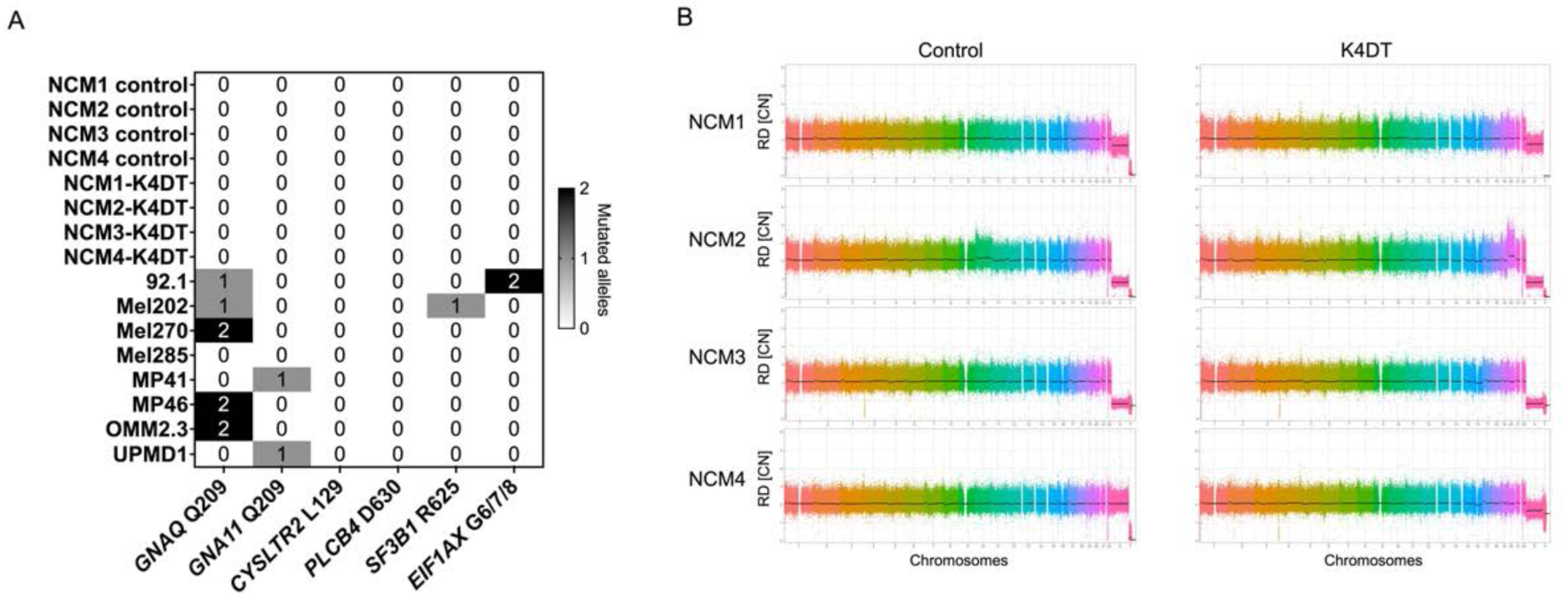
NCM-K4DT lines retain diploid copy number profiles and lack UM driver mutations. (A) Hotspot mutation map of key UM driver genes across cell panels. Targeted genotyping of *GNAQ*/*GNA11* Q209, *CYSLTR2* L129, *PLCB4* D630, *SF3B1* R625, and *EF1AX* G6/7/8 is shown for primary melanocytes (NCM controls) and immortalized derivatives (NCM-K4DT lines), with established UM cell lines included as references (i.e., 92.1, Mel202, Mel270, Mel285, MP41, MP46, OMM2.3, UPMD1). Shading and digit indicate the number of mutant-allele detection (grey (1) = heterozygous mutant; dark (2) = homozygous mutant; blank (0) = wild-type). (B) Genome-wide copy number (RD[CN]) profiles for all four donors before (Control) and after immortalization (K4DT).

In addition to driver gene mutations, UM frequently harbors copy number variations in chromosomes 3, 6 and 8 that carry significant clinical implications.^35^ Given that immortalization with SV40 or other viral oncogenes can introduce chromosomal instability,^18^ we next assessed whether our immortalized cell lines maintained genomic integrity. Genome-wide copy-number profiling confirmed a globally diploid karyotype in all NCM donors and derived K4DT lines, with no evidence of large-scale chromosomal gains or losses (Fig. 5B). We noted a subtle structural amplification on chromosome 20 in the NCM2-K4DT line, which likely reflects a focal copy number change acquired during immortalization rather than frank genomic instability, as it was not accompanied by additional widespread abnormalities.^36,37^ As expected from donor sex, NCM1 and NCM4 lines displayed a XX pattern, whereas NCM3 showed an XY configuration. In NCM2, a loss of the Y chromosome was observed (Fig. 5B). This is noteworthy because sexual chromosomal aberrations, such as loss of Y chromosome and derivative X chromosome, have been reported during immortalization and prolonged passaging of hTERT- or K4DT-immortalized human cells, as well as long-term cultured primary cells.^38,39^

### In vivo non-tumorigenicity of NCM-K4DT lines

To determine whether K4DT immortalization conferred tumorigenic potential, we performed an *in vivo* tumorigenicity assay using subcutaneous xenografts in immunodeficient mice. First, Luc2-tdTomato reporter expression was validated across the four NCM-K4DT lines using immunofluorescence and flow cytometry, confirming a transduction efficiency over 93% (Supplementary Fig. S3A). Then, a strong linear correlation between cell number and luminescent signal was demonstrated by *in vitro* luciferase assays (Supplementary Fig. S3B). We also used 92.1 cells as a positive control based on previous reports of tumor formation with this cell line.^40^ As expected, mice injected with 92.1-Luc2-tdTomato cells developed pronounced masses within 2 weeks post-injection and reached the experimental endpoint at 4-5 weeks (Fig. 6A). In contrast, mice injected with NCM-K4DT-Luc2-tdTomato lines developed small, measurable masses at 1-2 weeks post-injection that were no longer palpable by week 3 (Fig. 6B). *Ex vivo* analysis confirmed these findings, as mice injected with 92.1 cells developed large, firm tumors within the first 4-5 weeks, whereas mice injected with NCM-K4DT lines showed only small masses after 12 weeks (Fig. 6C). The recovered tumors from 92.1 were darkly pigmented with strong fluorescent and bioluminescent signals, whereas the masses from NCM-K4DT-injected mice were flat, fat-like Matrigel^®^ plugs with light pigmentation and weak but detectable fluorescent and bioluminescent signals, indicating that cells remained viable but non-tumorigenic after 12 weeks *in vivo* (Fig. 6D; Supplementary Fig. S3C-D). Taken together, these *in vivo* tumorigenic assays indicated that NCM-K4DT cells did not exhibit neoplastic growth potential, further reinforcing their suitability as non-transformed models.

**Figure 6.**
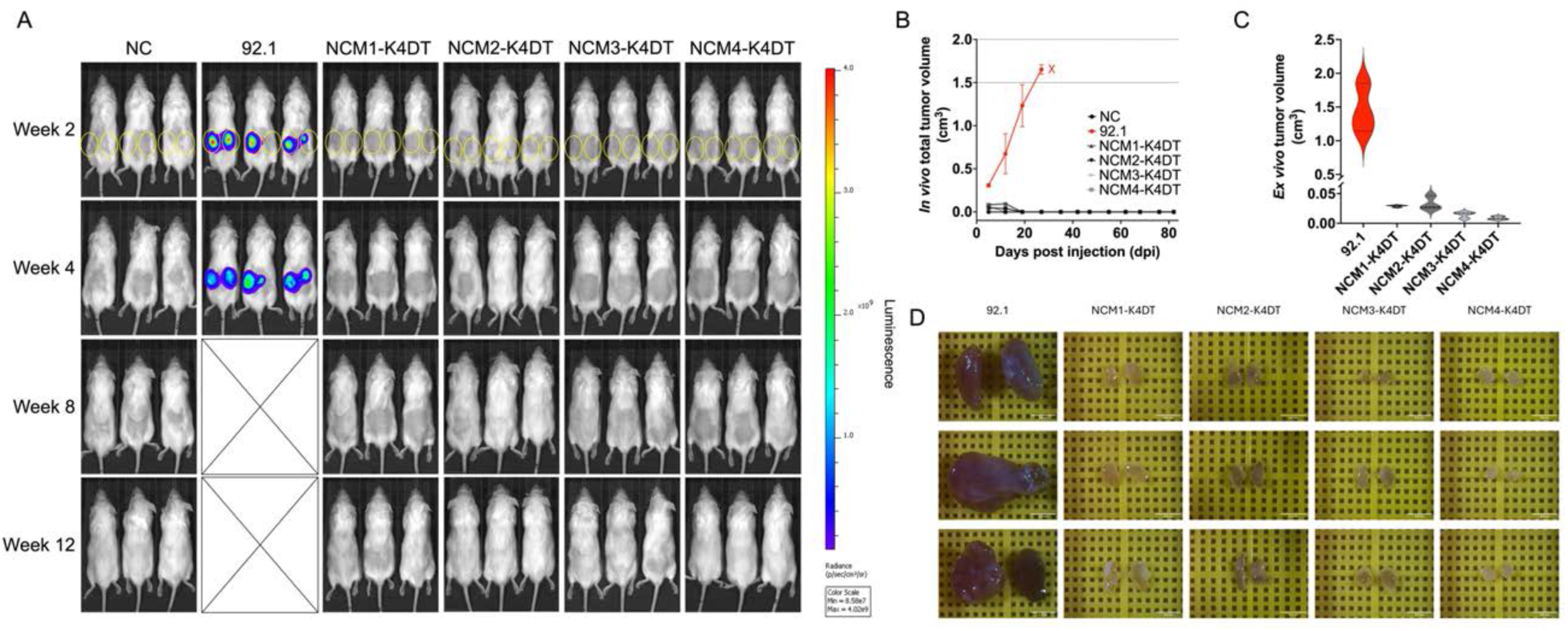
NCM-K4DT lines are non-tumorigenic *in vivo.* (A) Bioluminescence imaging (BLI) after subcutaneous injections in immunodeficient mice. Columns show groups (NC: PBS/Matrigel^®^ negative control; 92.1: positive control; four NCM-K4DT biological replicates); rows show images taken after 2, 4, 8 and 12 weeks post-injection. The color scale indicates radiance (photons s⁻¹ cm⁻² sr⁻¹); empty rectangles marked with an X means that all 3 mice in the 92.1 group were euthanized at week 4 because they had reached the limit point for tumor burden. (B) Longitudinal *in vivo* tumor volume measurements (cm^3^; mean ± SEM; n=3/group). (C) End-point *ex vivo* tumor volume (cm^3^; n=3/group). (D) Tumor/mass aspect for 92.1 and NCM-K4DT groups; scale bars = 0.5 cm.

### Generation of isogenic UM–associated mutations in NCM-K4DT lines using CRISPR-based editing

Primary melanocytes are notoriously difficult to genetically manipulate due to their slow proliferation and poor transfection efficiency, which has limited the application of CRISPR gene editing in this cell type. Given the importance of gene editing tools for studying melanocyte biology or generating early tumor development and progression models, we next evaluated whether immortalization could overcome these technical challenges and enable efficient CRISPR-mediated gene editing in NCM-K4DT cells. To assess the suitability of NCM-K4DT lines for gene editing, we applied SCIPpay constructs carrying engineered initiating UM mutations in *GNAQ* (Q209L; Fig. 7A) or *GNA11* (Q209L) and a secondary mutation in *BAP1* (C91G) that results in loss of function and is associated with poor UM prognostic.^41–43^ Among the tested constructs, p*GNA11*^Q209L^-HA yielded the lowest transfection efficiency, whereas p*GNAQ*^Q209L^-HA and p*BAP1*^C91G^-HA achieved higher levels of cells expressing the SpCas9-EGFP across all cell lines (Fig. 7B). Transfection efficiency, as assessed by EGFP+ cells 24 hours after electroporation, was readily detectable but varied between constructs. The cell viability remained generally high (>93%), except for the NCM4 donor that showed reduced number of live cells (80.8%; Fig. 7C). Quantification of precise insertion using ICE analysis (Synthego) further demonstrated that the p*GNAQ*^Q209L^-HA construct achieved the highest percentage of mutation positive alleles compared with the other tested constructs (Fig. 7D). Sanger sequencing further confirmed the precise co-introduction of the intended *GNAQ* mutation and the silent PAM mutation at the target locus in multiple cell lines (Fig. 7E). This mutational pattern is consistent with our SCIP-based approach and is unlikely to result from spontaneous mutations. These findings establish the feasibility of plasmid-based CRISPR editing in NCM-K4DT lines and support their use as isogenic models to study the functional impact of UM–associated mutations, although optimization of some constructs may be needed.

**Figure 7.**
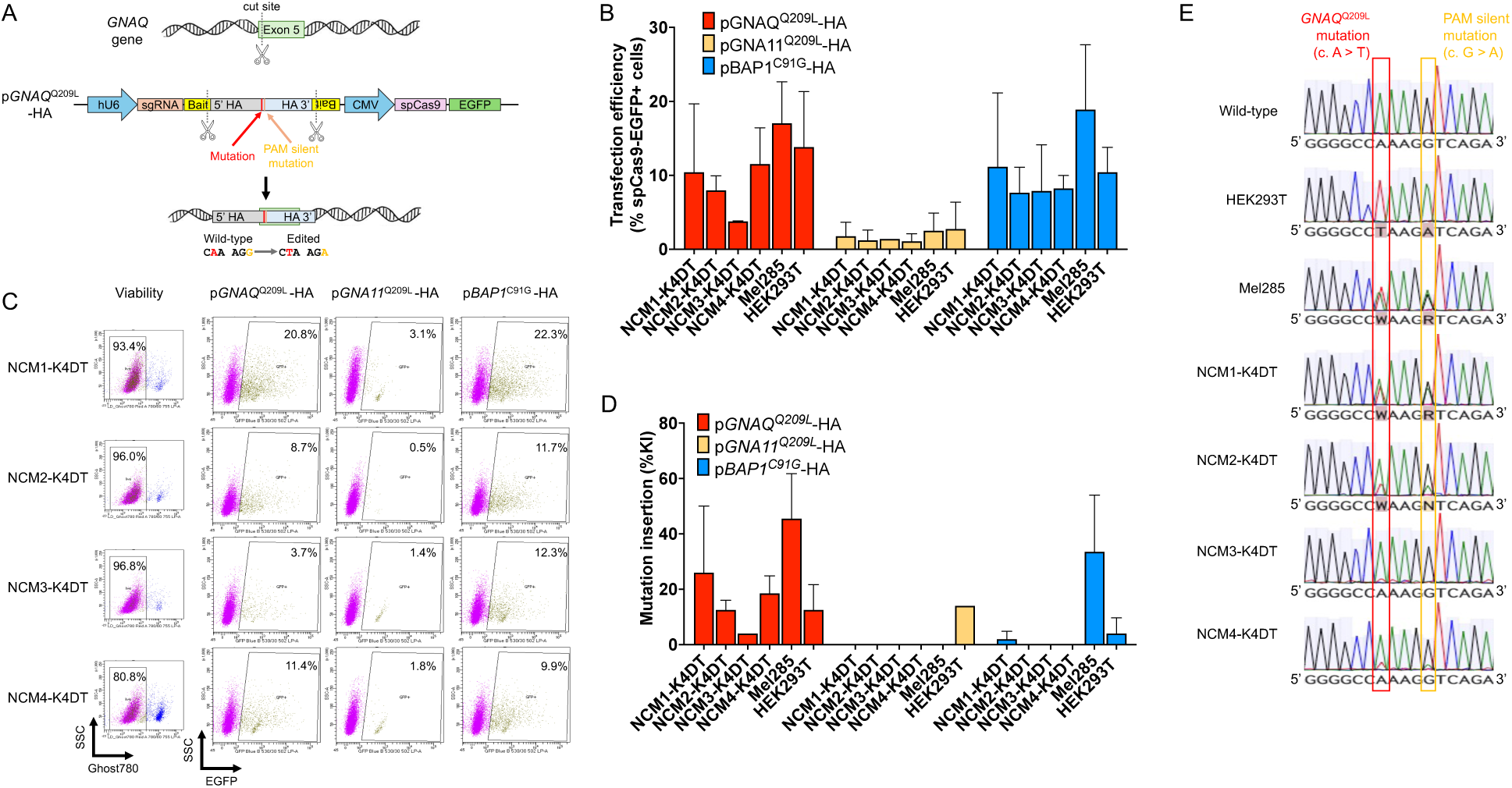
Efficiency of gene editing and targeted mutation insertion in NCM-K4DT lines. (A) Schematic of the SCIPpay construct designed to introduce the Q209L mutation into the human *GNAQ* gene. The donor plasmid contains the engineered mutation flanked by homology arms (HA) and the CRISPR complex with SpCas9-EGFP and gene-specific sgRNA. (B) Quantification of transfection efficiency (EGFP+ cells) across independent replicates. (C) Flow cytometry plots showing viability (Ghost Dye Red 780-) and transfection efficiency (EGFP+) in NCM-K4DT cells 24 hours after transfection with SCIPpay constructs targeting *GNAQ* (p*GNAQ*^Q209L^-HA), *GNA11* (p*GNA11*^Q209L^-HA), and *BAP1* (p*BAP1*^C91G^-HA) genes. Populations are indicated as percentages. (D) Quantification of mutation insertions at each target locus. (E) Sanger sequencing chromatograms confirming successful introduction of the intended mutations at the *GNAQ* locus in HEK293T, Mel285 and NCM-K4DT lines, compared with the wild-type sequence.

## Discussion

In this study, we established and characterized the first immortalized human choroidal melanocytes, addressing a need in ocular research models. While other ocular cell types, such as retinal pigment epithelium, are represented by commonly used immortalized lines (e.g., ARPE-19, hTERT-RPE1),^44,45^ research using choroidal melanocyte has relied exclusively on short-lived primary cultures.^1,6–15^ This dependence on primary NCMs imposes significant constraints, as their limited proliferative capacity restricts experimental throughput, and the requirement for multiple donors introduces genetic heterogeneity that confounds mechanistic studies. The NCM-K4DT model overcomes these limitations by providing a renewable, genetically tractable choroidal melanocyte system that preserves key lineage characteristics while remaining non-tumorigenic. This positions the NCM-K4DT model as a versatile platform for investigating both NCM biology and the early molecular events underlying UM.

Aside from choroidal melanocytes, the K4DT immortalization cassette is increasingly used to generate proliferative yet near-physiological models from diverse primary cells, including human epidermal melanocytes and ocular epithelial cells.^46,47^ This positions the NCM-K4DT model as a valuable addition to existing resources. The technique was originally described by Shinomi *et al.* in 2011 using human myogenic cells whereby the proliferative capacity of myocytes was increased, while preserving the myogenic phenotype, as well as retaining the potential to differentiate both *in vivo* and *in vitro*.^48^ Furthermore, compared to traditional immortalization techniques, the K4DT method resulted in minimal chromosomal instability, and substantially fewer differentially expressed cancer-related genes.^18,49^ Consistent with these observations, NCM-K4DT cells maintain core melanocytic lineage characteristics such as SOX-10 expression, as well as core melanin producing machinery. This includes melanosome-specific protein expression (PMEL, Melan-A), melanin producing enzymatic activity (TRP1 expression, dopachrome formation) and ultimately, melanin accumulation. Although it initially seemed as though melanin production was slightly decreased in immortalized cells, reducing growth-stimulating factors such as cholera toxin and PMA led to an accumulation of melanin in cells. This demonstrates that the reduced pigmentation in NCM-K4DT cells is consistent with dilution of melanin due to increased proliferation, rather than loss of melanocytic lineage identity. In this context, rapid cell division likely outpaces melanosome maturation and melanogenesis, resulting in progressive reduction of melanin content per cell.

The genetic tractability of NCM-K4DT lines makes them well-suited for introducing defined driver mutations to model cancer early events. To test the feasibility of introducing UM driver mutations in NCM-K4DT lines, we used the CRISPR-Cas9 technology to target three hotspot mutations: *GNAQ*^Q209L^, *GNA11*^Q209L^, and *BAP1*^C91G^. From a technical standpoint, gene editing in primary choroidal melanocytes is challenging due to their low transfection efficiency, and slow proliferation, which hinder the establishment of edited populations. While knock-in of *GNA11* and *BAP1* mutations was unsuccessful (likely due to construct-specific factors that could be addressed with optimization), we achieved mutational insertion of *GNAQ*^Q209L^. Importantly, we obtained efficient *GNAQ*^Q209L^ editing across all four donors at levels that were comparable to HEK293T cells. To our knowledge, this represents the first report of CRISPR-mediated knock-in of a recurrent UM driver mutation in human choroidal melanocytes. For this proof-of-concept study, edited populations were maintained as polyclonal pools rather than deriving single-cell clones, and we did not perform genome-wide off-target profiling or comprehensive pathway analyses. Nevertheless, this demonstrates the NCM-K4DT model is amenable to plasmid-based editing and homology-directed repair, thus enabling the generation of other isogenic cell lines to explore mutation-specific signaling dependencies in choroidal melanocytes and UM.

Although NCM-K4DT lines retain genomic stability and melanocytic identity, immortalization by enforced CDK4^R24C^/Cyclin D1/hTERT expression is unlikely to be completely neutral. Studies of hTERT-immortalized epithelial and fibroblast cells have shown that bypassing senescence is accompanied by discrete but detectable changes in DNA methylation and gene expression, particularly in pathways related to cell cycle control and differentiation, even when cells remain non-tumorigenic.^50,51^ Similarly, global transcriptomic comparisons indicated that K4DT-immortalized corneal epithelial cells cluster close to their primary counterparts but diverge for a restricted set of genes, consistent with mild epigenetic reprogramming associated with sustained proliferation.^49,52^ Nevertheless, by targeting the CDK4-Cyclin D1-RB axis downstream of mitogenic signaling but upstream of DNA replication and mitosis, this approach permits cell cycle entry while leaving multiple downstream checkpoints intact. This is evidenced by S/G2 accumulation, maintained genomic stability, and critically, non-tumorigenic behavior, since all NCM-K4DT lines did not form palpable masses after 12 weeks when grafted in flanks of immunodeficient mice.

The K4DT immortalization technique comprises two components that merit discussion regarding their potential impact on melanocyte biology. First, CDK4^R24C^ is a p16^INK4A^-resistant allele associated with hereditary cutaneous melanoma in humans,^53^ and other mutations in codon 24 (i.e., R24H, R24L) have been identified in somatic and germline cutaneous melanoma samples,^54^ as well as cell lines.^55^ However, in Cdk4^R24C^ knock-in mice, spontaneous melanoma development is rare and typically requires cooperating oncogenic stimuli such as HRAS^G12V^, indicating that this alteration alone is insufficient for transformation.^56,57^ The non-tumorigenic phenotype of NCM-K4DT lines supports this conclusion. Importantly, uveal and cutaneous melanocytes, and their corresponding malignancies, are biologically and pathologically distinct entities, characterized by different neural crest-derived developmental trajectories, oncogenic drivers, and signaling dependencies.^2,58–61^ Given these fundamental differences, CDK4^R24C^ expression does not preclude the use of NCM-K4DT lines for UM modeling. Nevertheless, investigators should exercise caution when designing experiments that directly target the CDK4/p16 axis, as the R24C mutation confers resistance to p16-mediated inhibition and would bypass such interventions. Next, the implication of Cyclin D1 overexpression must be considered given its association with UM. While elevated Cyclin D1 was observed in a subset of UMs and correlated with unfavorable outcomes,^62,63^ it is not a defining driver and does not arise from recurrent *CCND1* mutations or amplifications. Rather, increased Cyclin D1 in UM likely reflects convergent output from constitutively active upstream pathways, including PLCβ-PKC, MAPK, and Hippo-YAP signaling.^64^ In this context, Cyclin D1 enables cell cycle progression but does not actively drive oncogenic transformation.

The translational potential of the NCM-K4DT model lies in its versatility across choroid research applications. Our strategy balances the need for renewable cell supply with preservation of regulatory integrity. The retention of stress-responsive pathways, evidenced by elevated p21 expression, maintained genomic stability, and non-tumorigenic behavior, ensures that NCM-K4DT cells can respond to oncogenic perturbations without imposing a fully transformed baseline, making the model well-suited for studying signaling dependencies and therapeutic vulnerabilities in UM pathogenesis. Beyond cancer, the preservation of pigmentation capacity and lineage-specific markers supports studies of fundamental melanocyte functions, including melanogenesis, oxidative stress regulation, and intercellular interactions in the choroidal microenvironment. These pathways are not only central to UM but also play roles in non-neoplastic ocular disorders such as age-related macular degeneration, where the dysfunction of melanocytes and the retinal pigment epithelium contributes to the progression of the disease.^10,65,66^ Because aging is the strongest risk factor for cancer and most malignancies are diagnosed in late adulthood (median age at cancer diagnosis: 65-67 years),^67,68^ including UM which typically develops around 60 years of age,^69^ NCM-K4DT lines derived from both younger (∼48 years) and middle-aged (∼65 years) donors provide an attractive platform to investigate how age-associated changes in melanocytes and their microenvironment influence susceptibility to malignant transformation and treatment response.

In conclusion, to our knowledge, the NCM-K4DT lines constitute the first immortalized human choroidal melanocyte model, providing a renewable and biologically relevant platform for studying choroidal melanocyte homeostasis and malignant transformation. Their amenability to plasmid-based CRISPR editing and targeted introduction of UM-associated mutations further enables the generation of isogenic NCM–mutant pairs to dissect the functional impact of specific genetic lesions. By bridging the gap between short-lived primary cultures and tumor-derived models, they represent a critical resource for advancing our understanding of UM pathogenesis and for accelerating the development of targeted ocular therapies.

## Commercial Relationships Disclosure

A.F.R., None; A.M., None; V.G., None; K.C., None; A.D., None; S.L., None.

## Acknowledgments

The authors would like to thank Mrs. Julie Bérubé and Mr. Julien Blouin from Pr. Stéphanie Proulx’s Laboratory (Université Laval, Canada) for technical assistance with the isolation of choroidal melanocytes. We would also like to thank Dr. Darin Bloemberg (Novo Nordisk, Canada) for providing the pQCi backbone and for sharing his expertise on SCIP construct design. We also thank Pr. Sonia del Rincon (McGill University, Canada) for sharing the pCDH-EF1-Luc2-P2A-tdTomato transfer plasmid.

## Author contributions

A.F.R. designed and conducted the experiments and was responsible for writing the original draft of the manuscript and for editing. A.M. contributed to experiment design, execution, and manuscript editing. V.G. assisted with karyotyping analysis under the supervision of A.D. K.C. helped with NCM isolation from donor eyeballs and reviewed and edited the manuscript. S.L. provided conceptualization, preparation of the original draft of the manuscript, manuscript reviewing and editing, supervision, and funding acquisition. All authors have read and agreed to the submitted version of the manuscript.

## Supplementary data

**Supplementary Figure S1.**
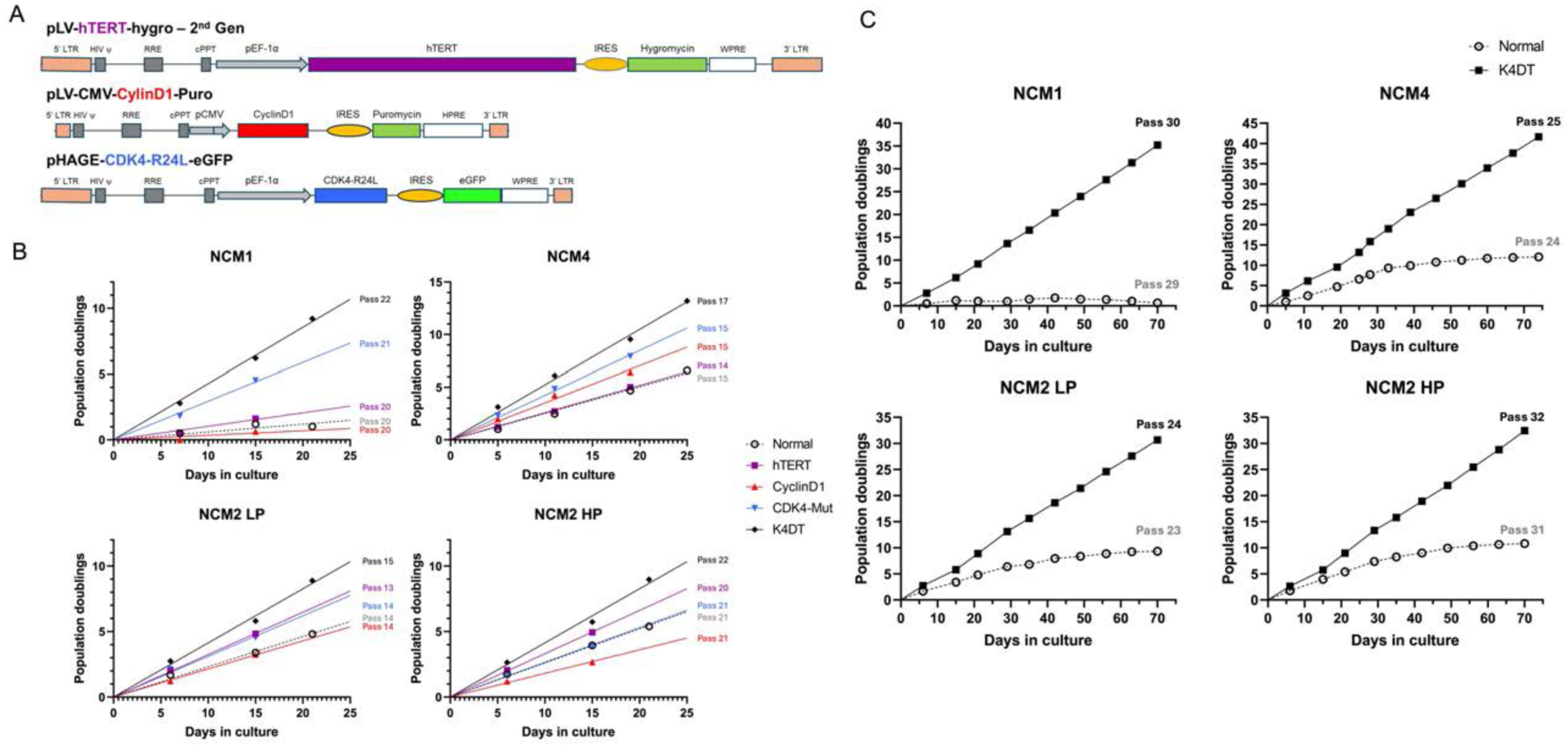
Pilot evaluation of hTERT, Cyclin D1, and CDK4-Mut transgenes on NCM proliferative capacity. (A) Schematic representation of lentiviral vectors used to express hTERT, Cyclin D1, or CDK4-Mut individually, as well as their combined expression (K4DT), in primary NCMs. (B) Short-term proliferation analysis showing population doublings of NCMs from four donors following transduction with individual transgenes (hTERT, Cyclin D1, or CDK4-Mut) or the combined K4DT construct. While individual transgenes did not consistently enhance proliferation relative to untransduced NCMs, combined K4DT expression resulted in increased population doublings across donors. Based on these pilot data, cultures expressing individual transgenes were discontinued after approximately three weeks. (C) Long-term growth curves comparing primary NCMs and K4DT-immortalized cells maintained in culture for up to 70 days. While the growth of primary NCM cultures plateaued, NCM-K4DT lines continued to undergo sustained population doublings over time.

**Supplementary Figure S2.**
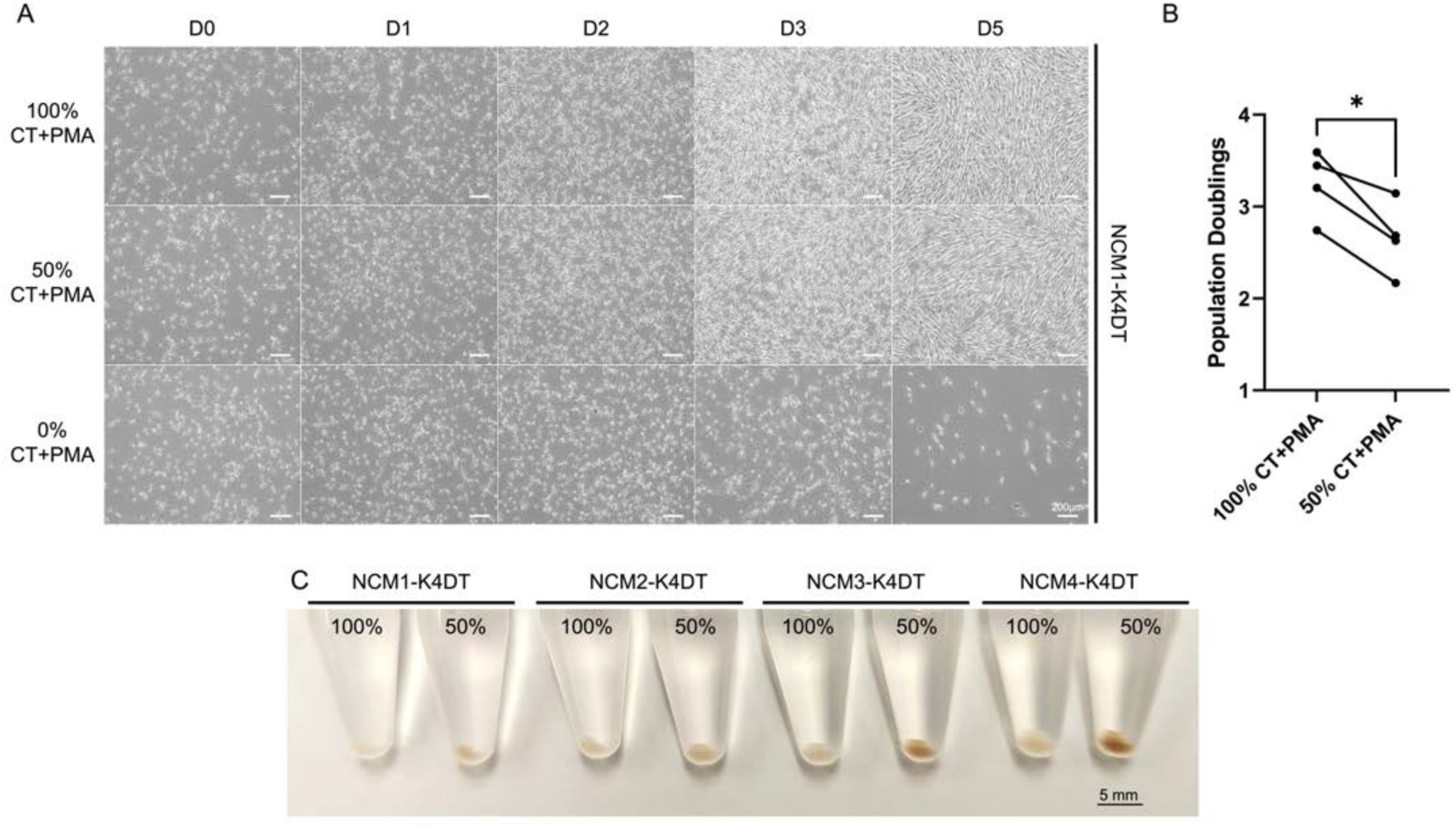
Reduced proliferation of NCM-K4DT lines is associated with increased melanin accumulation. (A) Phase-contrast images of NCM1-K4DT cells cultured in medium containing standard concentrations (100%) of cholera toxin (CT) and phorbol 12-myristate 13-acetate (PMA), reduced concentrations (50%), or no CT/PMA (0%). Over five days, cells cultured without CT and PMA were not viable, whereas cells maintained in 50% CT/PMA continued to proliferate, albeit more slowly than cells cultured under standard conditions. This trend was observed across all four NCM-K4DT donors (data not shown). (B) Quantification of population doublings for all NCM-K4DT lines after four days in culture under 100% or 50% CT/PMA conditions. Cells cultured in reduced CT/PMA exhibited a significant decrease in population doublings compared to standard conditions (*p* = 0.0179), consistent with a reduced proliferation rate. (C) Representative cell pellets (1.5×10⁶ cells) from NCM-K4DT lines cultured for four days in medium containing 100% or 50% CT/PMA. Cells cultured under reduced growth-stimulatory conditions displayed visibly darker pellets, consistent with increased melanin accumulation.

**Supplementary Figure S3.**
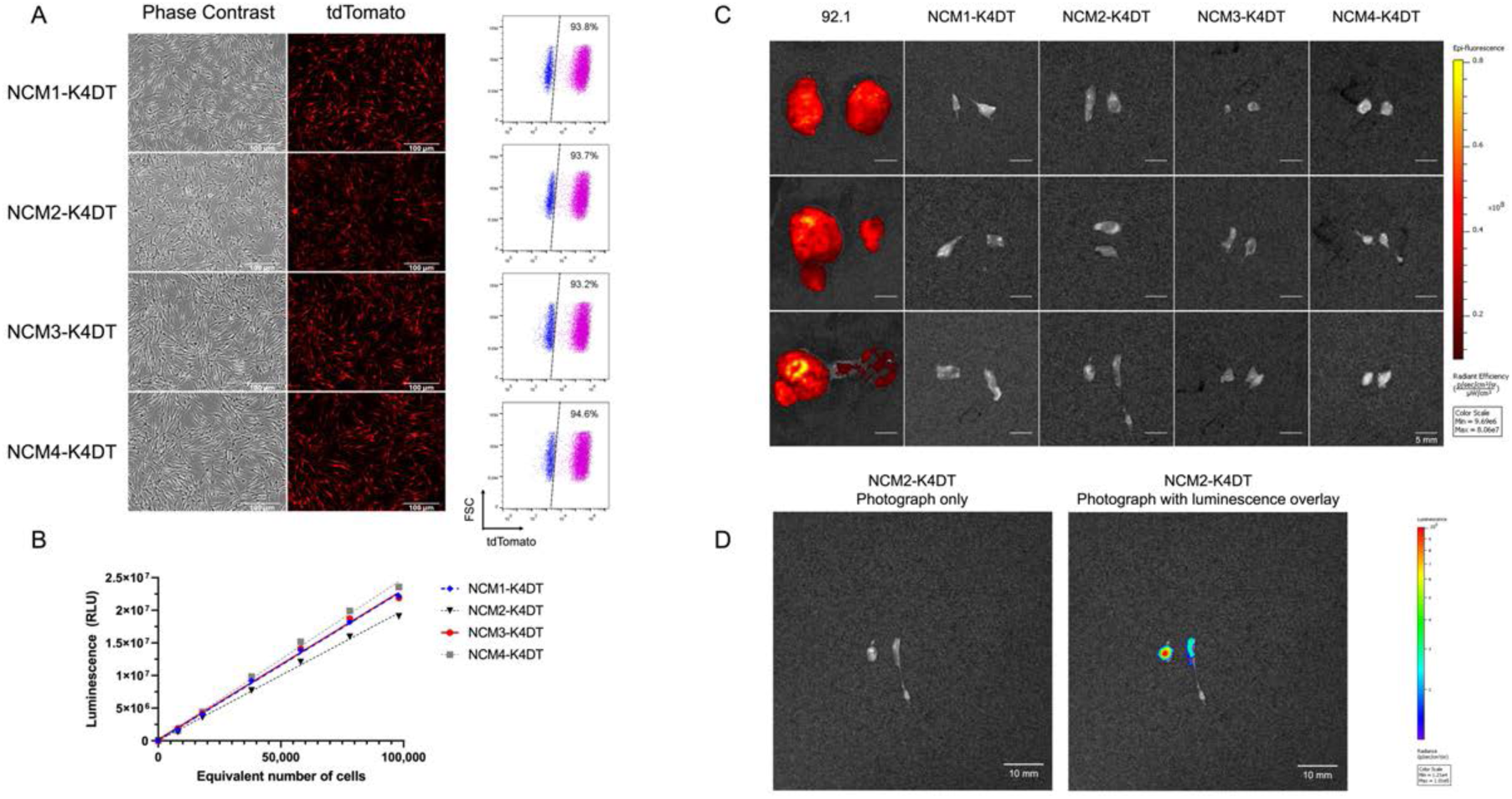
Generation and validation of Luc2-tdTomato-expressing NCM-K4DT lines. (A) Phase-contrast and fluorescence images demonstrating robust tdTomato expression across the four generated NCM-K4DT lines (left panel). Flow cytometric analysis confirming tdTomato expression in >93% of cells for all NCM-K4DT lines (right panel). (B) *In vitro* luciferase activity of Luc2-tdTomato-expressing NCM-K4DT lines prior to *in vivo* studies. Luminescence was measured across seven increasing cell numbers to generate standard curves, revealing a strong linear relationship between cell number and the luminescent signal, consistent with stable luciferase expression. (C) *Ex vivo* images of tumors and Matrigel^®^ plugs with fluorescence overlay. While strong tdTomato fluorescence was detected in tumors derived from the UM cell line 92.1, the fluorescence from NCM-K4DT Matrigel^®^ plugs was not detectable above background autofluorescence. (D) *Ex vivo* bioluminescence imaging of NCM2-K4DT Matrigel^®^ plugs performed immediately after euthanasia to assess cell viability and luciferase expression. Despite the absence of tumor formation, luminescent signal confirms the persistence of viable NCM2-K4DT cells 12 weeks post-injection. Due to variability in luciferin diffusion following intraperitoneal injection and the delay inherent to tissue dissection, luminescent signals could not be reliably captured at peak intensity; therefore, quantitative luminescent comparisons were not performed.

